# Targeted sequencing of mutations via RNA-templated gap filling of oligonucleotides for single-cell RNA-seq

**DOI:** 10.64898/2026.04.10.717677

**Authors:** Mirca Saurty-Seerunghen, Hower Lee, Martha Holdar, Kellie Wise, Michael Roach, Sara Moein, Ted Kang, Taobo Hu, Mats Nilsson, Luciano Martelotto, Anna Nam, Marco Grillo

**Affiliations:** Department of Pathology and Laboratory Medicine, Weill Cornell Medicine, New York, NY 10065, USA; Science for Life Laboratory, Department of Biochemistry and Biophysics, Stockholm University, Solna, 17165, Sweden; University of Adelaide, Adelaide Centre for Epigenetics, SAiGENCI, Adelaide, SA 5005, Australia; South Australian Immunogenomics Cancer Institute, University of Adelaide, Adelaide, SA 5005, Australia; Sandra and Edward Meyer Cancer Center, New York, NY 10065, USA

**Keywords:** Gap-filling, Single cell, RNA-seq, Mutation detection

## Abstract

We describe a method for targeted sequencing of expressed mutations in single cells. The approach exploits the reverse transcriptase and nick translation activities of the Bst DNA polymerase (Full Length) from *Bacillus stearothermophilus* (BstFL) to perform an RNA-templated gap filling and ligation between flanking probes. Integrated with probe-based single cell RNAseq workflows, it enables simultaneous whole transcriptomic and multiplexed mutation profiling from the same cell, overcoming key limitations of mutation detection in existing scRNA-seq approaches.

## Main

Detecting mutations on expressed mRNAs in single cell RNA-seq (scRNAseq) assays enables direct genotype-to-phenotype analysis in cancer and other somatically acquired diseases(1–4). However, detecting mutations in scRNA-seq is inherently difficult as scRNAseq coverage is sparse. We and others have developed methods to genotype in scRNA-seq that rely on capture of transcripts 5’ or 3’ ends; however, sensitivity is limited by the distance between the mutation and the captured transcript terminus(1–4). Genotyping methods based on plate-based full-transcript sequencing are constrained in their throughput to hundreds to a few thousands of cells(2, 3, 5, 6). Probe-based scRNAseq assays, in which pairs of oligonucleotides anneal adjacently on target mRNAs and are ligated to form sequencing-ready products (RASLseq, 10x Flex)(7, 8), offer improved sensitivity and experimental reliability. We previously described GoT-Multi, an RNA-templated ligation-based approach for mutation detection in single cells (9), but it presents challenges. The target variants must be known in advance to design allele-specific probes, and variable ligase specificity combined with probe competition may limit performance at loci with complex mutational landscapes.

To bypass these constraints, we designed an assay that exploits the RNA-templated DNA synthesis activity of BstFL polymerase to fill and define the gap sequence, converting unknown mutations into a ligation-competent substrate without requiring allele-specific probe design. In the first phase, Bst acts as a reverse transcriptase(10), extending a probe upstream of the mutation toward and across the gap. Spurious ligation is prevented by a 5’-OH on the terminus of the downstream probe (rather than the requisite 5’-phosphate), rendering it ligation-incompetent (Figure 1A). Once the gap is filled (Figure 1B-C), Bst cleaves off one or several nucleotides from the 5’ end via nick translation, exposing a 5’-phosphate end that makes the probe ligation-competent (Figure 1D-E). An RNA-templated DNA ligase (SplintR) then seals the nick(11) (Figure 1F). We validated this mechanism using an in situ assay targeting the 18s ribosomal RNA, employing padlock probes and rolling circle amplification (RCA) followed by Fluorescent in situ hybridization (FISH, Figure 1G-I)(12, 13). Critically, incorporating exonuclease-resistant backbone modifications into the downstream probe fine-tunes the competition between Bst’s polymerase and nick-translation activities, substantially improving overall gap-filling efficiency (Extended Data Fig. 1, Table S1).

**Fig. 1.**
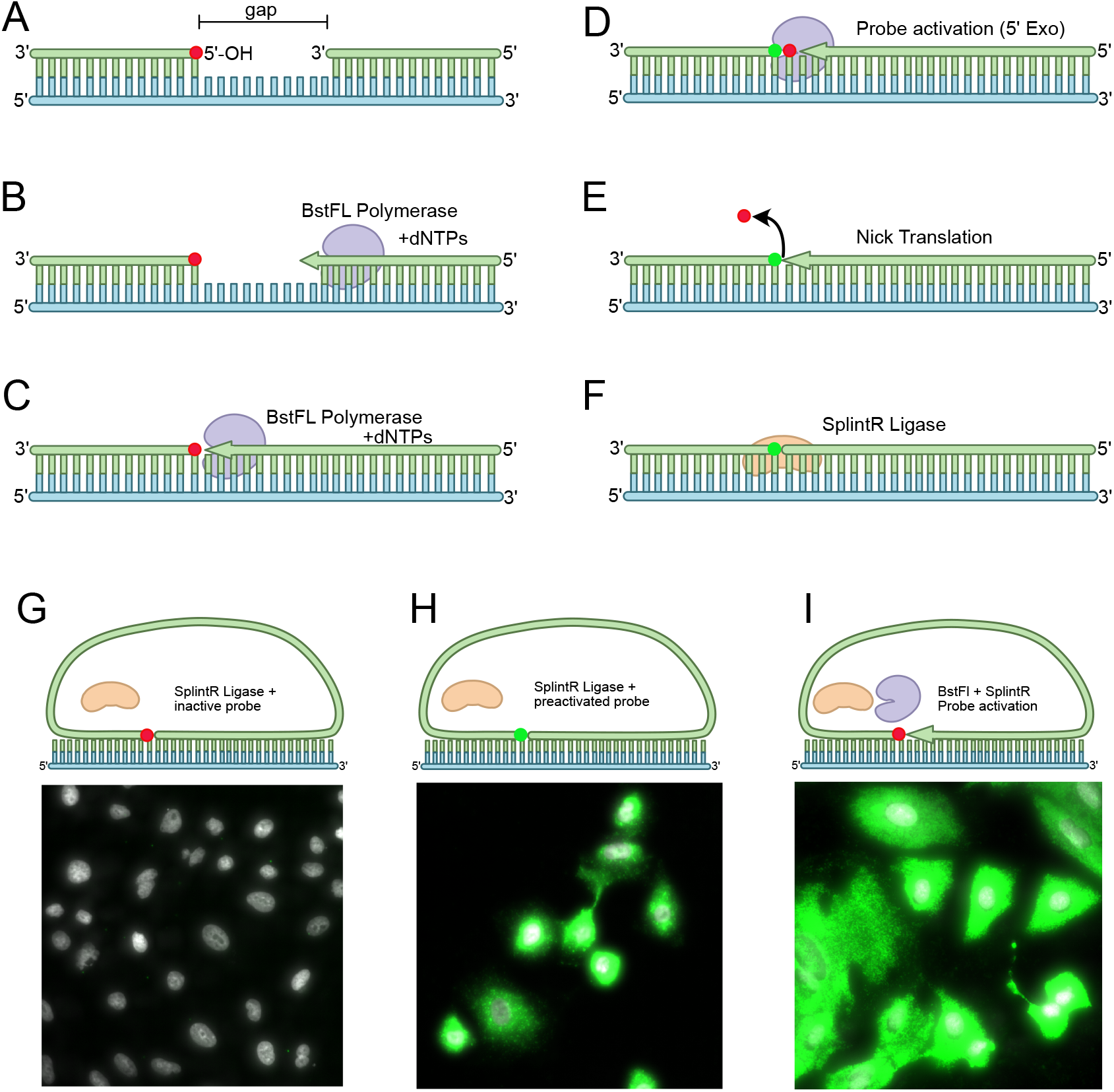
Molecular mechanism of an RNA-templated gap filling reaction: Panels A-F. Description of the molecular events involved in the RNA-templated gap filling reaction. A. Two probes flanking the mutation locus of interest are hybridized to the target region. The downstream probe (green DNA molecule left of the gap) has a terminal 5’OH nucleotide that makes it ligation-incompetent (red circle). B-C. The reverse-transcription activity of BstFL extends the 3’ end of the upstream probe (right) towards the downstream probe up to the complete filling of the gap. D. The nick translation activity of BstFL cleaves the 5’OH terminus, releasing the 5’OH terminal nucleotide (red circle) and exposing the 5’P of the following base (green circle). At this stage the downstream probe becomes ligation-competent. E. At the same time, the 3’ of the upstream probe grows into the nick, replacing the cleaved base. F. An RNA-templated DNA ligase (SplintR) efficiently seals the nick. Panels G-I. Validation of the gap-filling reaction chemistry with an in-situ assay. In the three panels, a padlock probe is hybridized against the 18s rRNA, then incubated with a reaction mix, rolling-circle amplified and finally incubated with a fluorescent probe to detect the rolling circle products (green signal, if present). The sequence of the probe is maintained identical in the 3 assays, and the cells are counterstained with DAPI G. The assay is performed using a 5’OH terminated padlock and incubated with SplintR ligase only. This serves as a negative control, confirming the ligation-incompetence of the padlock. H. The assay is performed using a 5’P terminated padlock and incubated with SplintR ligase. In this case the probe is ligation-competent, and serves a positive control for the efficient detection of the target RNA I. The assay is performed using a ligation-incompetent probe incubated with BstFL and SplintR. The target RNA can be detected only if nick translation happens, where the 5’OH is digested, the 5’P of the next nucleotide is exposed and the resulting gap is filled from the 3’end. The abundant green signal indicates successful activation of the probe via nick-translation.

We next designed GoT-Multi-Gap, an implementation of the Bst gap-filling assay in probe-based scRNAseq (Flex, 10x Genomics). Three cell lines, MCF-7(ER+ breast), SK-BR-3 (HER2+ breast) and LnCAP (prostate), were profiled by bulk RNAseq to identify 61 cell-line-specific single nucleotide variants (SNV), including polymorphisms, as a ground truth. Flanking probes were designed for each SNV across a range of gap sizes and mutation positions within the gap (Table S2). Cells were mixed in equal proportions and subjected to the gap-filling assay (Figure 2A; Materials and Methods), followed by the 10x Flex V1 workflow, enabling simultaneous gene expression and multiplexed mutation analysis from a single experiment. Importantly, the gap-filling step did not detectably perturb the downstream 10x Genomics’ Flex assay performance. Comparisons to the standard Flex experiment on the same cell lines showed similar unique transcript (UMI) counts and gene counts, and strong gene-wise correlations (Extended Data Fig. 2A-B). The three cell line populations remain fully distinguishable (Figure 2B) by canonical marker gene expression (Figure 2C), and differentially expressed markers (Extended Data Fig. 1C). We targeted 64 loci, including three loci with no variants as negative controls (Extended Data Fig. 3A-B). Loci with no variants displayed expected wildtype sequences with minimal background noise, likely due to sequencing or PCR errors (Extended Data Fig. 3A). Of the 61 targeted loci, 11 displayed less than 3 genotyped cells, 8 of which could be explained by low gene expression (Extended Data Fig. 3B). Among the targets with at least 10 genotyped cells and 3 mutant cells (38 targets), mutant calls displayed a high degree of specificity to the expected cell line(s) (range: 80-100%, Figure 2D). We systematically evaluated variables affecting mutation capture efficiency (Extended Data Fig. 3B). Target mRNA abundance was the main contributing factor: normalizing gap-filling capture against gene expression levels (normalized for the number of probes per transcript), revealed a moderate positive correlation (Figure 2E, Pearson’s R = 0.37, P < 0.01), consistent with reduced detection at lowly expressed loci. Within the recommended 44-72% GC content window for each probe half(14), flanking probe GC content had no detectable effect on efficiency; outside this range, sensitivity dropped significantly (Figure 2F, Extended Data Fig. 3C). Gap length showed no correlation with efficiency after normalizing for gene expression levels (Figure 2G, top panel). Longer gaps were less efficiently filled under controlled conditions in the FISH assay (Extended Data Fig. 1D-E), but this effect was not recovered in the single-cell data, likely due to sequence heterogeneity across targets masking gap length-dependent effects. Mutation position within the gap had no detectable effect on capture efficiency, with no bias toward variants proximal to either flanking probe (Figure 2G, bottom panel). The assay therefore captures variants across the full gap length with comparable sensitivity, with target abundance representing the primary deterministic variable identified in this analysis.

**Fig. 2.**
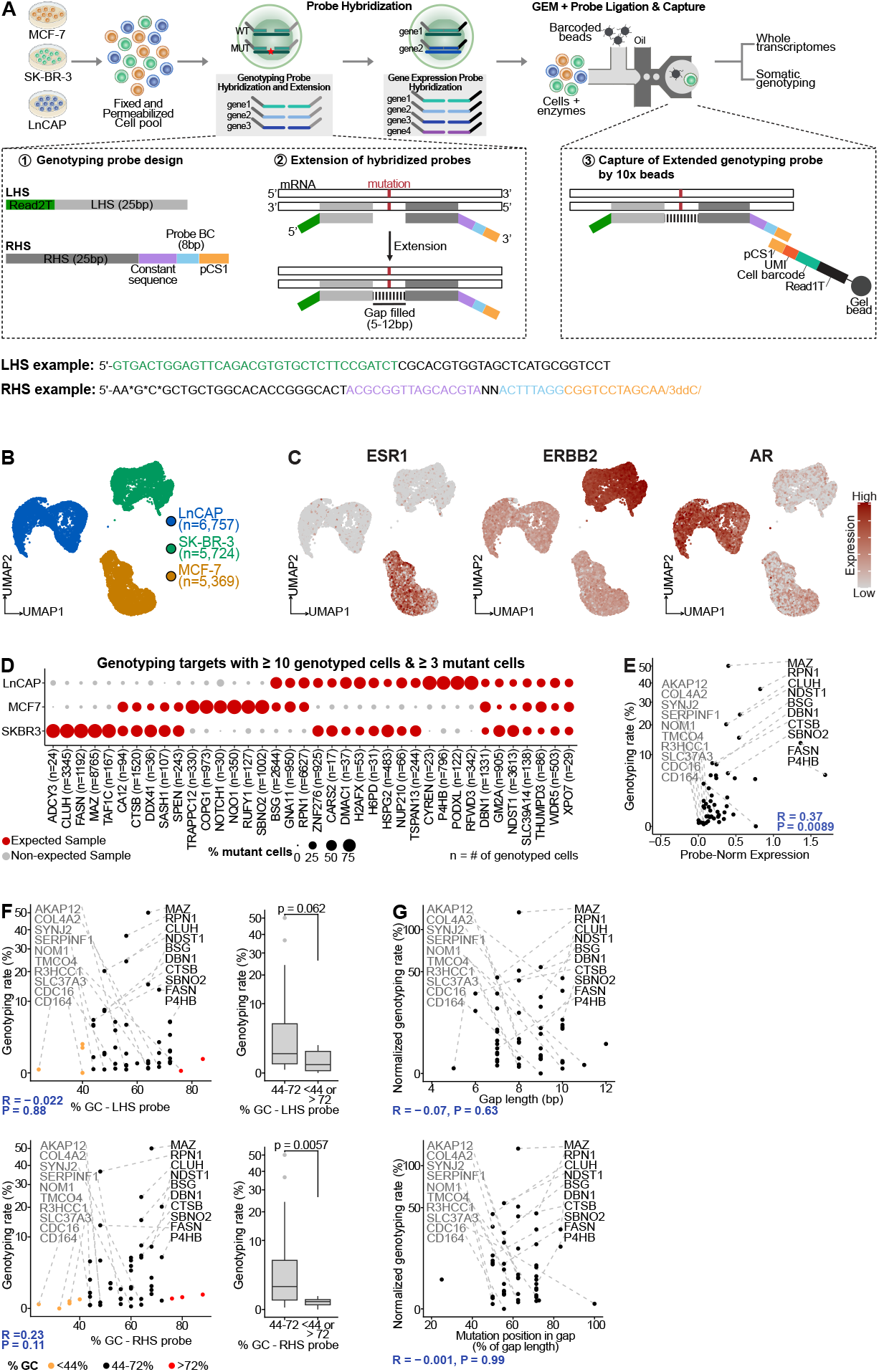
Genotyping from single-cell transcriptomes via RNA-templated gap filling: A. Schematic of experimental workflow. Cells from three cell lines are pooled, fixed and permeabilized. They are then hybridized with genotyping probes targeting specific mutation loci. Bottom-left insert shows the Left-Hand Side (LHS) and Right-Hand Side (RHS) probe design. The LHS probe has two components: a 25 bp sequence upstream of the mutation and a Read2T handle (green). The RHS probe consists of four components: (1) a 25 bp sequence downstream of the mutation on the 5’ end, (2) a constant sequence (purple), (3) a 8 bp probe barcode (blue), and (4) the probe capture sequence (pCS1; orange) recognized by the gel bead. The 5’ end of the RHS probe is initially ligation-incompetent (5’OH). The LHS and RHS probes hybridize to mRNA molecules of the corresponding gene. Bst FL polymerase extends the sequences from the 3’ end of the LHS to fill the gap to the 5’ end of the RHS probe, then nicks the RHS 5’OH terminal nucleotide and undergoes controlled nick translation and ligation. The ligated probe pair can then be captured by the 10x Genomics gel bead just like the standard gene expression (GEX) probes. Separate libraries are generated for gene expression and genotyping probes. The LHS and RHS probes for one representative mutation is provided as an example. The same colors as in the probe design are used in the example. B. Uniform manifold approximation and projection (UMAP) overlaid with cell line identity. SK-BR-3 and MCF-7 cells are human HER2+ and HER2-ER+ breast cancer cell lines, respectively. LnCAP is a human AR+ prostate cancer cell line. C. UMAP of cells from the three cell lines (n = 17,850 cells), overlaid with expression levels of ESR1 (ER), ERRB2 (HER2) and AR. D. Dot plot showing percentage of mutant cells in each cell line for all targets. E. Genotyping rate of each target (n=50) relative to mean expression in samples expected to be mutated (expected samples). Genotyping rate corresponds to the percentage of genotyped cells in the expected samples. Expression values were normalized by the number of GEX probes from 10x Genomics. Top 10 most or least genotyped targets are highlighted in black and grey, respectively. P-value from Pearson correlation analysis. F. Genotyping rate of each target (n=50) relative to the GC content of its LHS and RHS probes (top-left and bottom-left panels, respectively). Genotyping rate and targets highlighted as in Figure 2E. P-value from Pearson correlation analysis. Box plots comparing genotyping rate of targets with probes that have acceptable GC content (44-72%) or not (<44% or > 72%) (top-right and bottom-right panels). P-value from Wilcoxon test. G. Genotyping rate of each target (n=50) relative to the length of the gap being filled (left panel) or position of mutation in the gap (right panel). Genotyping rate normalized by mean expression in the expected samples. Mutation position in gap corresponds to the position of the mutation in the gap relative to gap length (% of gap length). Top 10 most or least genotyped targets are highlighted in black and grey, respectively. P-value from Pearson correlation analysis.

## Conclusions

In summary, we present an RNA-templated gap-filling strategy that enables mutation detection in single-cell transcriptomes. By leveraging the dual enzymatic activity of Bst polymerase, this approach overcomes key limitations of existing ligation-based assays. Mutation capture efficiency is robust across diverse sequence contexts, with target transcript abundance as the primary, and largely predictable, determinant of detection sensitivity. Importantly, integration with the 10x Genomics’ Flex workflow preserves transcriptomic integrity enabling simultaneous gene expression and multiplexed variant profiling, laying the groundwork for systematic and scalable interrogation of expressed genetic variation at single-cell resolution across the full transcriptome.

## ACKNOWLEDGEMENTS

We’d like to thank the members of the Nilsson lab and Daniel Gyllborg for constructive discussions, and the following funding sources: To MN: Project grants from Knut and Alice Wallenberg foundation (2024/0058), the Swedish Research Council (2024-02533), and Cancerfonden (24-3457)

## References

1. Nam, Anna S and Kim, Kyu-Tae and Chaligne, Ronan and Izzo, Franco and Ang, Chelston and Taylor, Justin and Myers, Robert M and Abu-Zeinah, Ghaith and Brand, Ryan and Omans, Nathaniel D and Alonso, Alicia and Sheridan, Caroline and Mariani, Marisa and Dai, Xiaoguang and Harrington, Eoghan and Pastore, Alessandro and Cubillos-Ruiz, Juan R and Tam, Wayne and Hoffman, Ronald and Rabadan, Raul and Scandura, Joseph M and Abdel-Wahab, Omar and Smibert, Peter and Landau, Dan A. Somatic mutations and cell identity linked by Genotyping of Transcriptomes. Nature 571, 355–360 (2019).

2. Giustacchini, Alice and Thongjuea, Supat and Barkas, Nikolaos and Woll, Petter S and Povinelli, Benjamin J and Booth, Christopher A G and Sopp, Paul and Norfo, Ruggiero and Rodriguez-Meira, Alba and Ashley, Neil and Jamieson, Lauren and Vyas, Paresh and Anderson, Kristina and Segerstolpe, Åsa and Qian, Hong and Olsson-Strömberg, Ulla and Mustjoki, Satu and Sandberg, Rickard and Jacobsen, Sten Eirik W and Mead, Adam J. Single-cell transcriptomics uncovers distinct molecular signatures of stem cells in chronic myeloid leukemia. Nat. Med. 23, 692–702 (2017).

3. Rodriguez-Meira, Alba and Buck, Gemma and Clark, Sally-Ann and Povinelli, Benjamin J and Alcolea, Veronica and Louka, Eleni and McGowan, Simon and Hamblin, Angela and Sousos, Nikolaos and Barkas, Nikolaos and Giustacchini, Alice and Psaila, Bethan and Jacobsen, Sten Eirik W and Thongjuea, Supat and Mead, Adam J. Unravelling intratumoral heterogeneity through high-sensitivity single-cell mutational analysis and parallel RNA sequencing. Mol. Cell 73, 1292–1305.e8 (2019).

4. van Galen, Peter and Hovestadt, Volker and Wadsworth, Ii, Marc H and Hughes, Travis K and Griffin, Gabriel K and Battaglia, Sofia and Verga, Julia A and Stephansky, Jason and Pastika, Timothy J and Lombardi Story, Jennifer and Pinkus, Geraldine S and Pozdnyakova, Olga and Galinsky, Ilene and Stone, Richard M and Graubert, Timothy A and Shalek, Alex K and Aster, Jon C and Lane, Andrew A and Bernstein, Bradley E. Single-cell RNA-seq reveals AML hierarchies relevant to disease progression and immunity. Cell 176, 1265–1281.e24 (2019).

5. Picelli, Simone and Björklund, Åsa K and Faridani, Omid R and Sagasser, Sven and Winberg, Gösta and Sandberg, Rickard. Smart-seq2 for sensitive full-length transcriptome profiling in single cells. Nat. Methods 10, 1096–1098 (2013).

6. Picelli, Simone and Faridani, Omid R and Björklund, Asa K and Winberg, Gösta and Sagasser, Sven and Sandberg, Rickard. Full-length RNA-seq from single cells using Smart-seq2. Nat. Protoc. 9, 171–181 (2014).

7. Li, Hairi and Qiu, Jinsong and Fu, Xiang-Dong. RASL-seq for massively parallel and quantitative analysis of gene expression. Curr. Protoc. Mol. Biol. Chapter 4, Unit 4.13.1–9 (2012).

8. 10x Genomics. Chromium fixed RNA profiling reagent kits for singleplexed samples. Tech. Rep., Pleasanton, CA (2024).

9. Pak, Minwoo and Saurty-Seerunghen, Mirca S and Wise, Kellie and Abera, Tsega-Ab and Lama, Chhiring and Parghi, Neelang and Kang, Ted and Sun, Xiaotian and Gao, Qi and Bao, Liming and Roshal, Mikhail and Allan, John N and Furman, Richard R and Martelotto, Luciano G and Nam, Anna S. Co-mapping clonal and transcriptional heterogeneity in somatic evolution via GoT-Multi. Cell Genom. 6, 101036 (2026).

10. Shi, Chao and Shen, Xiaotong and Niu, Shuyan and Ma, Cuiping. Innate reverse transcriptase activity of DNA polymerase for isothermal RNA direct detection. J. Am. Chem. Soc. 137, 13804–13806 (2015).

11. Lohman, Gregory J S and Zhang, Yinhua and Zhelkovsky, Alexander M and Cantor, Eric J and Evans, Thomas C. Efficient DNA ligation in DNA–RNA hybrid helices by Chlorella virus DNA ligase. Nucleic Acids Res. 42, 1831–1844 (2014).

12. Larsson, Chatarina and Koch, Jørn and Nygren, Anders and Janssen, George and Raap, Anton K and Landegren, Ulf and Nilsson, Mats. In situ genotyping individual DNA molecules by target-primed rolling-circle amplification of padlock probes. Nat. Methods 1, 227–232 (2004).

13. Krzywkowski, Tomasz and Hauling, Thomas and Nilsson, Mats. In situ single-molecule RNA genotyping using padlock probes and rolling circle amplification. Methods Mol. Biol. 1492, 59–76 (2017).

14. 10x Genomics. Custom probe design for gem-x flex v2. Tech. Rep.

