## Extended data and Methods for "Targeted sequencing of mutations via RNA-templated gap filling of oligonucleotides for single-cell RNA-seq"

**Supplementary Information**

**Mirca Saurty-Seerunghen[1 a], Hower Lee[2 a], Martha Holdar[2], Kellie Wise[3,4], Michael Roach[3,4], Sara Moein[1], Ted Kang[1], Taobo Hu[2], Mats Nilsson[2 *], Luciano Martelotto[3,4 *], Anna Nam[1,5 *], and Marco Grillo[2 *]**

[1]Department of Pathology and Laboratory Medicine, Weill Cornell Medicine, New York, NY 10065, USA
[2]Science for Life Laboratory, Department of Biochemistry and Biophysics, Stockholm University, Solna, 17165, Sweden
[3]University of Adelaide, Adelaide Centre for Epigenetics, SAiGENCI, Adelaide, SA 5005, Australia
[4]South Australian Immunogenomics Cancer Institute, University of Adelaide, Adelaide, SA 5005, Australia
[5]Sandra and Edward Meyer Cancer Center, New York, NY 10065, USA

**This PDF contains the extended methods, supplementary tables, extended data figures, and supplementary references.**

### Extended Data and Methods

**Cell cultures.** MCF-7 and SK-BR-3 are cancer cell lines originating from a female patient with breast adenocarcinoma aged 69 and 43 years old, respectively while LnCAP is a cancer cell line originating from a male patient with metastatic prostate carcinoma aged 50 years old. MCF-7 cells were cultured in Dulbecco's Modified Eagle Medium (DMEM; Gibco, #11965-092) supplemented with 10% fetal bovine serum (FBS, Gibco, #16000-044) and 2 mM L-glutamine (Gibco, #25030-081). SK-BR-3 cells were maintained in McCoy's 5A Modified Medium (Thermo Fisher Scientific, #16600082) supplemented with 10% FBS (Gibco, #16000-044). LnCAP cells were cultured in RPMI-1640 (Gibco, #11875-093) supplemented with 10% FBS (Gibco, #16000-044). All cell lines were cultured at 37 °C in a humidified atmosphere with 5% $CO_2$. Cells were dissociated using TrypLE™ Express Enzyme (Thermo Fisher Scientific, #12604013) and cryopreserved in freezing medium containing 90% FBS and 10% dimethyl sulfoxide (DMSO; Merck, #41639). Cells were frozen at a density of 1 × 106 cells/mL in a CoolCell container (Corning) at –80 °C overnight, then transferred to –150 °C for long-term storage. Cell viability and concentration were assessed pre- and post-freezing using the Luna-FX7 Automated Cell Counter (Logos Biosystems) with the AO/PI Viability Kit (#F23011). Post-thawing, viability consistently exceeded 85%. On the day of fixation, cells were thawed in a 37 °C water bath and washed twice with PBS (without $Ca^{2+}$/$Mg^{2+}$) containing 0.05% BSA (Miltenyi Biotec, #130-091-376). A459 cells (lung cancer cell line) were cultured in DMEM (Gibco) without phenol red and l-glutamine, supplemented with 10% FBS (Sigma), 2 mM l-glutamine (Sigma) and 1× PEST (Sigma). To prepare cell samples, we treated confluent cell lines with 0.25% (w/v) trypsin-EDTA (Sigma) and resuspended them in culturing medium. Resuspended cells were then seeded on Superfrost Plus slides (Thermo) placed in a 150 mm × 25 mm Petri dish (Corning), and culturing medium was added to a final volume of 25 ml. Three milliliters of resuspended cells were used to seed five slides. Cells were incubated in the same previous conditions 12–24 hours before fixation.

**In-situ assay** Padlock probes targeting the human 18s rRNA were designed with a variable gap size of 0, 10, 20 nucleotides. The probes were ordered as DNA ultramers from Integrated DNA Technologies. As a positive control, we phosphorylated an aliquot of the 18s 0-gap padlock probe, incubating it under the following conditions: 6 pmol/µL of padlock probes were incubated with 0.2 U/µL of PNK (NEB# M0201SL) and 1 mM ATP in a reaction volume of 25 µL at 37 °C for 2 hours. The reaction was inactivated at 65 °C for 20 minutes. For all of the other experiments, the probes were left non-phosphorylated. A549 cells were seeded on SuperFrost slides and fixed in 3% formaldehyde for 30 minutes at room temperature, then stored at -80 °C until needed. The slides were thawed and allowed to reach room temperature, then progressively dehydrated in ethanol 70% for 2 minutes and ethanol 100% for 2 minutes. Slides were air-dried and a 50 µL Secure Seal chamber (#SA8R, Grace Biolabs) was applied on the slide. Cells were washed twice with PBS and then incubated overnight at 45 °C with 2x SSC, 20% Ethylene Carbonate (#E26258-100G, Sigma Aldrich) and 10 nM padlock probe. The next day the hybridization mix was removed and the slides were washed 3 times with 10% formamide in 2x SSC at room temperature, followed by 2 washes in PBST (PBS 1x + 0.1% Tween). For non-phosphorylated probes, the slides were then incubated at 45 °C for 2 hours with 1x ThermoPol buffer (#B9004S, NEB), 2 mM $MnCl^2$, 500 µM dNTPs (#N0447L, NEB), 1 U/µL of RiboProtect Hu (#RT35L, Qiagen/Blirt), and 0.25 U/µL of BstFL polymerase (#M0328S, NEB). For pre-phosphorylated probes, the slides were incubated for 2 hours in PBS 1x at 45 °C. All the slides were washed twice with PBST and then incubated with a ligation mix containing 0.05 µM of RCA primer, 1X T4 RNA Ligase Buffer (#B0216L, NEB), 0.5 mM $MnCl^2$, 10 µM ATP, 1U/µL RiboProtect Hu and 0.25 U/µL SplintR ligase (#M0375L, NEB) at 37 °C for 2 hours. The slides were washed twice with PBST and a Rolling Circle

DRAFT

Amplification mix containing 1x Phi29 Buffer, 5% Glycerol, 0.25 mM dNTPs, 0.2 µg/mL Recombinant Albumin (#B9200L, NEB) and 1 U/µL Phi29 Polymerase (#4002, Montserate Biotechnology) was incubated at 30 °C overnight. The slides were washed twice with PBST incubated for 30 minutes at RT with a detection mix containing 2xSSC, 20% formamide, 0.1 µM fluorescently-labeled probe and 0.5 µg/µl DAPI. The slides were finally washed twice with PBST, mounted with SlowFade Gold (#S36937, ThermoFisher) and imaged under a wide-field microscope as previously described(1).

**Selection of SNPs** Candidate Single-Nucleotide Polymorphisms (SNP) for the LOH panel were selected using a custom R-based workflow developed in-house. Publicly available variant annotation resources were used to identify informative coding SNPs with valid dbSNP rs identifiers, and candidate loci were prioritized for intermediate population allele frequencies to maximize heterozygosity and LOH informativeness. Population allele frequencies, including overall AF as well as AFR, AMR, EAS, EUR, and SAS subgroup frequencies, were annotated using the Bioconductor packages SNPlocs.Hsapiens.dbSNP155.GRCh38 and MafDb.1Kgenomes.phase3.GRCh38. Gene annotation and chromosomal band information were retrieved from Ensembl using biomaRt. For assay design, genomic sequence context surrounding each candidate SNP was extracted from the hg38 reference genome, and reverse-complement sequences were generated for variants located on the minus strand. Variants with unfavorable sequence characteristics, such as extended homopolymeric G/C stretches, were excluded. The annotated candidates were then reviewed to retain one informative SNP per gene while ensuring broad genome-wide representation. This process resulted in a final panel of 50 SNPs distributed across chromosomes 1-22 for downstream analysis.
Next, the SNPs were filtered to keep those on expressed genes and detected in one of the cell lines tested. SNPs were identified in each cell line by applying the rnavar workflow from the nf-core suite (Nextflow(2), v25.10.4) onto bulk RNA-sequencing data. The workflow was run with default parameters in a Singularity containerized environment on a high-performance computing system at Weill Cornell Medicine. In brief, sequencing reads from the FASTQ files were aligned to the GRCh38 reference genome and annotated. Germline SNPs and indels were called using the GATK4 HaplotypeCaller within the rnavar workflow. Known human genetic variants were identified by using the NCBI SNP database as reference (`All_20180418.vcf.gz` downloaded from `https://github.com/MRCIEU/vcf-reference-datasets/blob/master/get_dbsnp.sh`). The identified SNPs were then filtered to keep variants with QualByDepth (QD) $> 2$, StrandOddsRatio (SOR) $< 3$, RMSMappingQuality (MQ) $> 40$, MappingQualityRankSumTest (MQRankSum) $> -12.5$ and ReadPosRankSumTest (ReadPosRankSum) $> -8$, as per GATK guidelines. Of the 50 SNPs, 44 SNPs detected in at least one cell line were retained (Table S1). Three additional SNPs that are not detected in any of the cell lines (ASAP2 rs2715860, RAP1GAP2 rs55904912 and MOB3A rs2074894) were included to assess specificity of the GoT-Multi-Gap workflow.

**RNA-sequencing and somatic variant calling** Total RNA from LnCAP, MCF7, and SKBR3 cells was prepared using the RNA Easy Mini Kit (Qiagen) according to the manufacturer's instructions. RNA was submitted for library preparation (i.e., Illumina Stranded mRNA Prep) and sequencing to the Australian Genome Research Facility (AGRF). Sequencing was carried out as per their standard RNA Sequencing Service (`https://www.agrf.org.au/service-guides/#nextgenseq`).RNA-seq reads in FASTQ format were first aligned to the hg38 human reference genome using STAR(3) (v2.7.10) and sorted with SAMTools(4) (v1.17). Homozygous and heterozygous single nucleotide polymorphism (SNP) variants were called with VarScan(5) (v2.3.9). Variant calls were combined with gene annotations and filtered to only include variants within gene exons using BEDTools(6) (v2.30.0). Manual inspections of a priori marker variants were performed with Integrative Genomics Viewer(7) and further filtering visualisations of all variants were performed in R with tidyverse(8) (v2.0.0). As a result, seventeen somatic mutations present in genes that are expressed and in at least one cell line were retained and targeted in this study (Table S2).

**GoT-Multi-Gap experimental workflow** To simultaneously capture whole transcriptomic data and genotyping data, Genotyping of Transcriptomes via Gap filling (GoT-Multi-Gap) was performed by adapting the 10x Genomics Fixed RNA Profiling (FRP) platform, a probe-based method for mRNA profiling, to incorporate RNA-templated gap filling (Figure 2A). Cells were fixed following the Fixation of Cells & Nuclei for Chromium Fixed RNA Profiling kit (10x Genomics, #PN1000414). Cells were then first hybridized with custom genotyping probe pools for 20 hours at 42 °C, followed by 3 washes with 1x Post-hybridization buffer (10x Genomics, #PN2000533). The custom genotyping probes correspond to probes designed to target specific mutation loci identified from RNA-sequencing data and validated using databases for single nucleotide polymorphisms (SNP). For each mutation, two probes were included (Figure 2A, Table S1): (1) a 25 bp left-hand side probe complementary to the mutation region a few bases upstream of the mutation site on 5'-3' strand (LHS), and (2) a 25 bp right-hand side probe containing the sequence a few bases downstream of the mutation. The mutation site must be in between the LHS and RHS probes, with a gap of maximum 5-12 bp between the probes. The LHS probe has two components: a 25 bp sequence upstream of the mutation and a Read2T handle (Figure 2A). The RHS probe consists of four components: (1) a 25 bp sequence downstream of the mutation on the 5' end, (2) a constant sequence, (3) a 8 bp probe barcode, and (4) the probe capture sequence (pCS1) recognized by the gel bead (Figure 2A). The 5' end of the RHS probe is initially ligation-incompetent (5'OH). The LHS and RHS probes hybridize to mRNA molecules of the corresponding gene. Bst FL polymerase extends the sequences from the 3' end of the LHS to fill the gap to the 5' end of the RHS probe, then nicks the RHS 5'OH terminal nucleotide and undergoes controlled nick translation and ligation. The ligated probe pair can then be captured by the 10x Genomics gel bead similar to the standard gene expression probes. Each genotyping probe pool contained probes for multiple mutations. Genotyping probe pairs for each mutation were ordered separately from Integrated DNA Technologies (IDT), resuspended at 100 µM and then pooled together before hybridization. To distinguish the genotyping from the gene expression reads and specifically amplify the genotyping ones, the genotyping probes were designed with a different PCR handle (Read2T handle: 5'-GTGACTGGAGTTCAGACGTGTGCTCTTCCGATCT) compared to the gene expression probes (Figure 2A, Table S1). After genotyping probe hybridization and washing, cells are subjected to the Gap Filling reaction (1x ThermoPol, 2 mM MnCl2, 500 µM dNTP, 1 U/µL RiboProtect and 1 0.25 U/µL Bst FL) and incubated for 3 hours at 45 °C. After gap filling, cells were washed twice with 2x PBS-T and and once with 0.5x PBS + 0.02% BSA. Following, cells were hybridized with 10x Genomics gene expression probes (Human WTA Probes BC001, #PN2000495) for 20 hours at 42 °C following Chromium Single Cell Gene Expression Flex user guide(9). Next, cells were washed 3 times with 1x Post-hybridization buffer (10x Genomics, #PN2000533) to remove unbound probes, counted and then loaded onto Chromium Next GEM Chip Q. The GEM incubation and recovery, the pre-amplification step and library construction steps were performed as per manufacturer's instructions. The transcriptomic profile was obtained from the gene expression library whereas the cell surface protein library contained the genotyping libraries. The genotyping libraries were spiked into the gene expression libraries to be sequenced together on a NovaSeq 6000 (Illumina, San Diego, CA). The cycle settings were as follows: 150 cycles for Read 1, 150 cycles for Read 2, 10 cycles for i7 sample index and 10 cycles for i5 sample index.

**GoT-Multi-Gap scRNA-seq data processing** Gene expression libraries were processed using Cell Ranger (v7.1.0) with the cellranger-multi pipeline, aligned to the human genome GRCh38 with default parameters. Downstream analysis was performed using Seurat(10) (v4.3.0) with default parameters unless otherwise specified. Cells with more than 20% mitochondrial gene percentage, less than 100 genes and UMI counts > or < 3 standard deviations from the mean UMI were filtered out (Extended Data Fig. 2A). The gene expression matrix was normalized, and the top 2000 variable genes selected by variance stabilizing transformation. Expression data was scaled and principal component analysis (PCA) was performed, retaining 30 principal components for downstream embedding. Cells were visualized using UMAP and clustered using graph-based Louvain clustering. Cluster identity was assigned manually based on differentially expressed genes identified withFindAllMarkers (Wilcoxon rank-sum test, minimum cell fraction 10%, top 2000 variable genes), cross-referenced against known canonical markers for each cell line (Figure 2B-C, Extended Data Fig. 2C). Doublets were predicted using scDblFinder(11) (v1.12.0) and DoubletFinder(12) (v2.0.3). To reduce false positives, only cells flagged as doublets by both tools and displaying mixed marker expression were excluded from downstream analysis.

**10x Genomics Flex scRNA-seq data processing** To assess whether the addition of custom probes and the gap filling step could affect gene expression probe binding, we compared our data against a standard 10x Genomics Flex experiment on the same cell lines, previously published by Pitino et al(13). Gene expression reads from that dataset were downsampled to match the reads number per cell of the current study, then aligned to the human genome GRCh38 using Cell Ranger (v7.0.0) with the cellranger-multi pipeline (default parameters) and processed as described above.

**Genotype calling using IronThrone pipeline** Genotyping library was performed using the IronThrone pipeline(14, 15). A modified version supporting multiple mutation targets simultaneously was used (https://github.com/AnnaNamLab/GoT-Multi-ML). To ensure correct probe hybridization and extension (gap filling), genotyping amplicon reads were first screened for the expected LHS sequence, and 10 bp of the RHS sequence, designated as 'primer' and 'shared' sequences in the IronThrone pipeline. Reads passing this filter were then processed through IronThrone to identify wild-type or mutant sequences at the expected position, as previously described(15). Since the genotyping FASTQ files contain the reads for multiple mutation targets, reads were first subsetted by expected extended probe sequence (allowing up to 20% mismatches) prior to IronThrone processing, improving computational efficiency. For each mutation, cells with sufficient genotyping reads ($\geq$ 2 reads per UMI) and at least one mutant UMI were called 'mutant'; cells with sufficient reads but no mutant UMI were called 'wildtype'; all remaining cells were classified as 'unprofiled'. Genotyping rate corresponds to the percentage of cells from the cell line(s) expected to be mutated (hereafter designated as 'expected sample') that have been profiled for a given target relative to the total number of cells from that cell line(s). Genotyping specificity corresponds to the percentage of mutant cells in the expected sample relative to the total number of mutant cells. Of the 61 SNVs targeted (Extended Data Fig. 3C), eleven were not detected. Low target transcript abundance accounted for eight of them., and also explains reduced genotyping rates at loci such as CD164 or AKAP12. Low probe GC content represents an additional contributing factor for a subset of undetected targets (e.g., CCT4).

**Quantification and statistical analysis** Statistical details are reported in the corresponding figure legends. Unless otherwise specified, p-values were adjusted using Benjamini-Hochberg (BH) FDR-correction, with statistical significance defined as an adjusted p-value < 0.05. Exceptions are noted in the figure legends or Methods section.

**Extended data and figures** Table S1. Padlock probes sequences for the in situ-assay

Table S2. Probe sequences for each mutation targeted in this study.

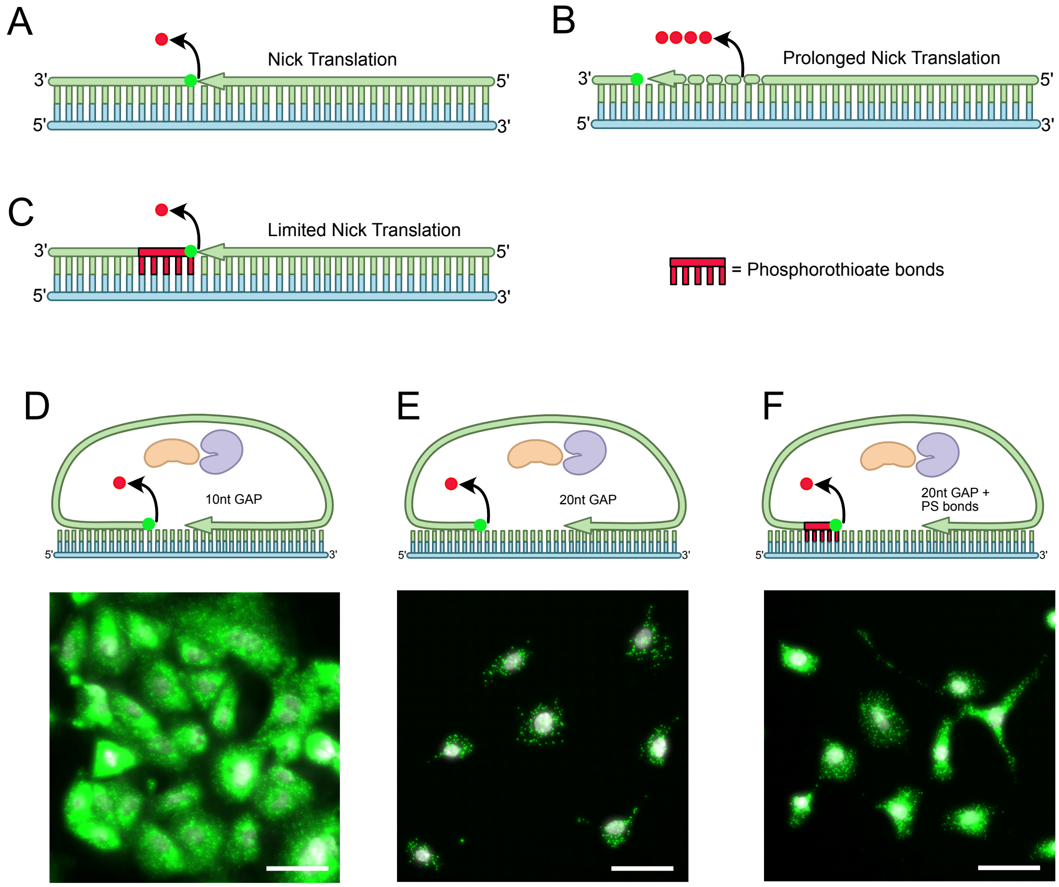


**Fig. Extended Data 1. Optimization of the gap filling efficiency via limited nick translation.** A-C. Design of a strategy for limited nick-translation: A. The nick translation reaction removes the 5'OH terminal nucleotide from the downstream probe, making it competent for ligation. If the balance between the nicking and ligation rates is not optimal, or if the two steps are not performed simultaneously, nick translation can proceed to the next nucleotides. B. If nick translation proceeds at an excessive rate, Bst will digest multiple nucleotides from the downstream probe and fill them back from the upstream one. This might lead to an excessive shortening of the downstream probe, potentially reducing the efficiency of the assay. C. The nick translation activity can be limited to the first nucleotide by introducing a stretch of phosphorothioate bonds in the downstream oligo. This modification makes the oligonucleotide resistant to the exonuclease activity of Bst, preventing excessive digestion of the downstream probe. D-F. Testing the impact of gap length and restricted nick-translation activity on the efficiency assay. D. 18s rRNA labeling with a 10 nucleotide gap inactivated probe, via gap filling and probe activation. This serves as a reference efficiency for a very abundant target. The level of detection is somewhat reduced compared with Figure 1I (0nt gap), as expected as a consequence of incomplete gap filling efficiency. E. 18s rRNA labeling via gap filling and activation of a 20 nucleotide gap probe show further reduction in the detection efficiency, suggesting that either larger gaps are more difficult to efficiently fill, or that enzyme imbalances produce excessive digestion of the padlock probe. F. The low efficiency of assay E can be partially rescued by the introduction of PS bonds in the downstream arm of the padlock probe, suggesting that restricted nick translation promotes more efficient gap filling.

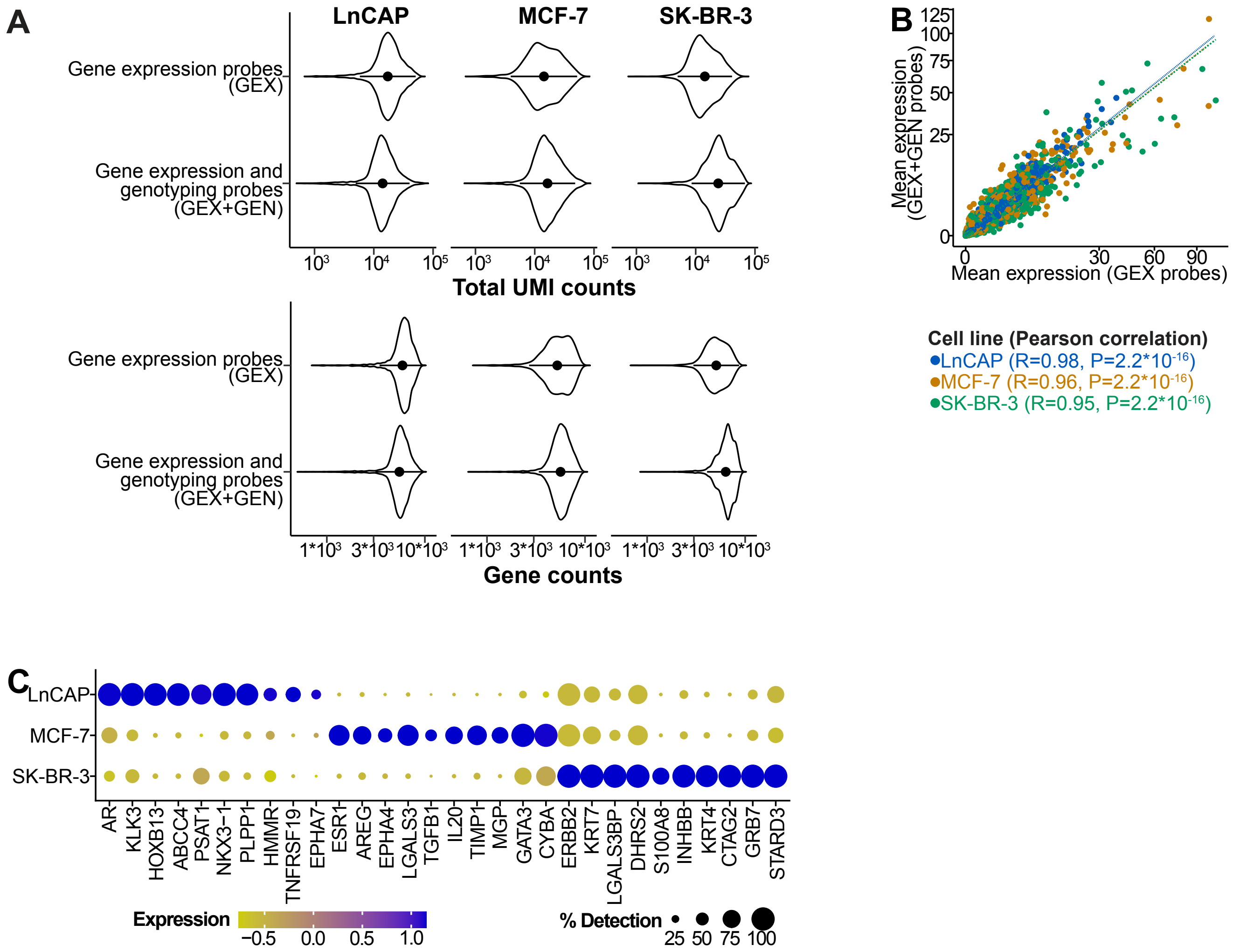


**Fig. Extended Data 2. Effect of gap filling on single-cell transcriptomes.** A. Violin plots showing number of UMIs (left panel), and genes (right panel) detected per cell in each cell line after filtering based on quality control (QC) metrics (methods). Cells from an experiment using regular FRP protocol with gene expression probes only1 (top) or from this study with gene expression and genotyping probes (bottom). B. Correlation plot showing expression levels of genes captured in this study relative to those detected when only gene expression probes are included. P-values from Pearson correlation analysis. C. Dot plot showing expression levels of genes characteristic of each cell line.

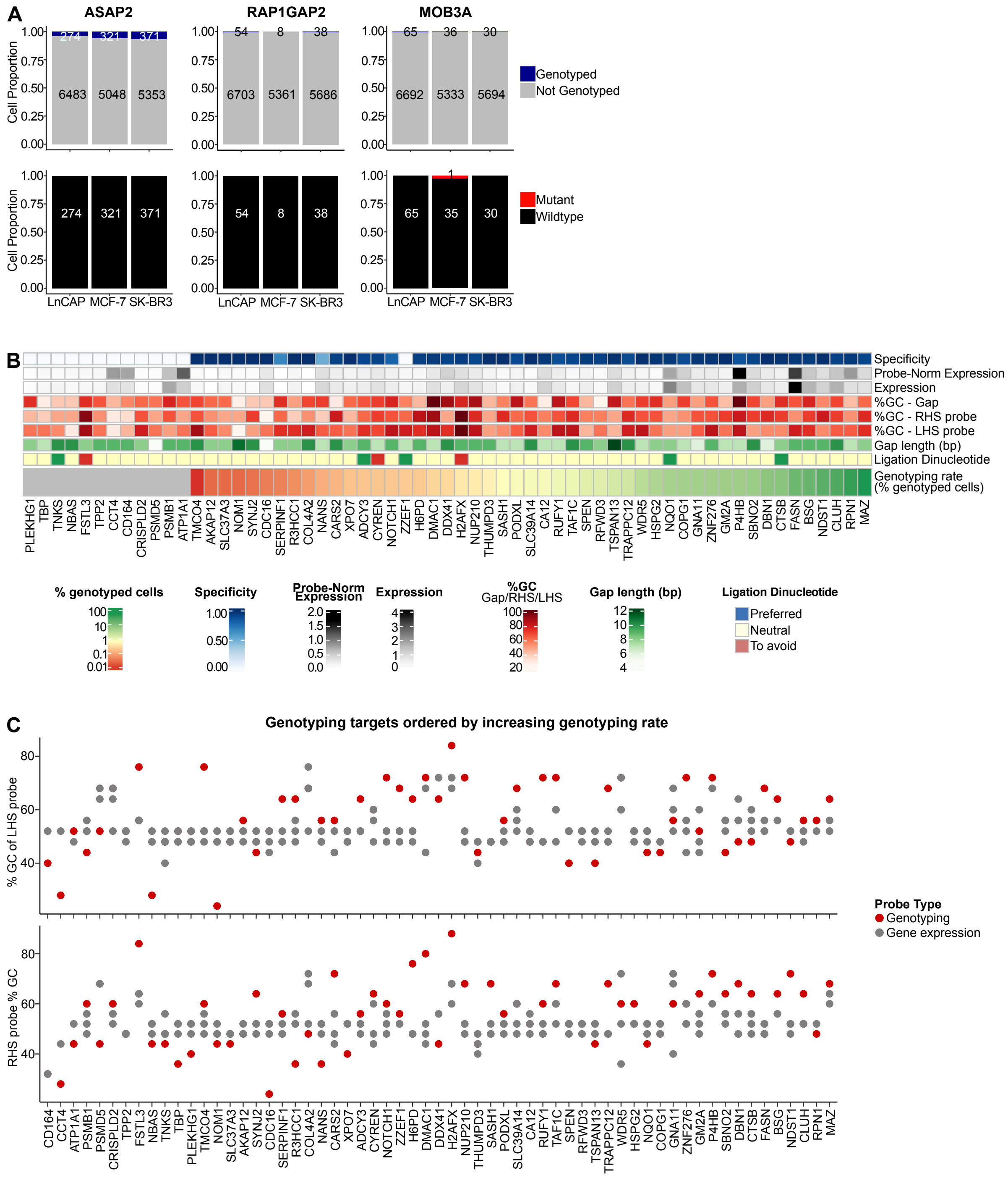

**Fig. Extended Data 3. Efficiency of genotyping from single-cell transcriptomes via RNA-templated gap filling.** A. Bar plots showing genotyping rate (top panels) or genotyping status (bottom panels) for three mutations not detected in any cell line (negative controls). B. Heatmap showing features of the genotyping probes for all mutations profiled in this study (n=61). Probe-Norm Expression corresponds to mean gene expression normalized by the number of GEX probes. Genotyping rate (% genotyped cells) corresponds to percentage of genotyped cells in the expected samples.C. Plot showing GC content of LHS (top) and RHS (bottom) sequences for gene expression (grey) or genotyping (red) probes for each target (n=61). Targets ranked by increasing genotyping rate. Genotyping rate corresponds to percentage of genotyped cells in the expected samples.

### Supplementary References


1. Gyllborg, Daniel and Langseth, Christoffer Mattsson and Qian, Xiaoyan and Choi, Eunkyoung and Salas, Sergio Marco and Hilscher, Markus M and Lein, Ed S and Nilsson, Mats. Hybridization-based in situ sequencing (HybISS) for spatially resolved transcriptomics in human and mouse brain tissue. *Nucleic Acids Res.* **48**, e112 (2020).
2. Di Tommaso, Paolo and Chatzou, Maria and Floden, Evan W and Barja, Pablo Prieto and Palumbo, Emilio and Notredame, Cedric. Nextflow enables reproducible computational workflows. *Nat. Biotechnol.* **35**, 316–319 (2017).
3. Dobin, Alexander and Davis, Carrie A and Schlesinger, Felix and Drenkow, Jorg and Zaleski, Chris and Jha, Sonali and Batut, Philippe and Chaisson, Mark and Gingeras, Thomas R. STAR: ultrafast universal RNA-seq aligner. *Bioinformatics* **29**, 15–21 (2013).
4. Danecek, Petr and Bonfield, James K and Liddle, Jennifer and Marshall, John and Ohan, Valeriu and Pollard, Martin O and Whitwham, Andrew and Keane, Thomas and McCarthy, Shane A and Davies, Robert M and Li, Heng. Twelve years of SAMtools and BCFtools. *Gigascience* **10**, giab008 (2021).
5. Koboldt, Daniel C and Chen, Ken and Wylie, Todd and Larson, David E and McLellan, Michael D and Mardis, Elaine R and Weinstock, George M and Wilson, Richard K and Ding, Li. VarScan: variant detection in massively parallel sequencing of individual and pooled samples. *Bioinformatics* **25**, 2283–2285 (2009).
6. Quinlan, Aaron R. BEDTools: The Swiss-army tool for genome feature analysis: BEDTools: The Swiss-army tool for genome feature analysis. *Curr. Protoc. Bioinformatics* **47**, 11.12.1–34 (2014).
7. Robinson, James T and Thorvaldsdóttir, Helga and Winckler, Wendy and Guttman, Mitchell and Lander, Eric S and Getz, Gad and Mesirov, Jill P. Integrative genomics viewer. *Nat. Biotechnol.* **29**, 24–26 (2011).
8. Wickham, Hadley and Averick, Mara and Bryan, Jennifer and Chang, Winston and McGowan, Lucy and François, Romain and Grolemund, Garrett and Hayes, Alex and Henry, Lionel and Hester, Jim and Kuhn, Max and Pedersen, Thomas and Miller, Evan and Bache, Stephan and Müller, Kirill and Ooms, Jeroen and Robinson, David and Seidel, Dana and Spinu, Vitalie and Takahashi, Kohske and Vaughan, Davis and Wilke, Claus and Woo, Kara and Yutani, Hiroaki. Welcome to the tidyverse. *J. Open Source Softw.* **4**, 1686 (2019).
9. 10x Genomics. Chromium fixed RNA profiling reagent kits for singleplexed samples. Tech. Rep., Pleasanton, CA (2024).
10. Hao, Yuhan and Hao, Stephanie and Andersen-Nissen, Erica and Mauck, 3rd, William M and Zheng, Shiwei and Butler, Andrew and Lee, Maddie J and Wilk, Aaron J and Darby, Charlotte and Zager, Michael and Hoffman, Paul and Stoeckius, Marlon and Papalexi, Efthymia and Mimitou, Eleni P and Jain, Jaison and Srivastava, Avi and Stuart, Tim and Fleming, Lamar M and Yeung, Bertrand and Rogers, Angela J and McElrath, Juliana M and Blish, Catherine A and Gottardo, Raphael and Smibert, Peter and Satija, Rahul. Integrated analysis of multimodal single-cell data. *Cell* **184**, 3573–3587.e29 (2021).
11. Germain, Pierre-Luc and Lun, Aaron and Garcia Meixide, Carlos and Macnair, Will and Robinson, Mark D. Doublet identification in single-cell sequencing data using scDblFinder. *F1000Res.* **10**, 979 (2021).
12. McGinnis, Christopher S and Murrow, Lyndsay M and Gartner, Zev J. DoubletFinder: Doublet detection in single-cell RNA sequencing data using artificial nearest neighbors. *Cell Syst.* **8**, 329–337.e4 (2019).
13. Pitino, Emanuele and Pascual-Reguant, Anna and Segato-Dezem, Felipe and Wise, Kellie and Salvador-Martinez, Irepan and Crowell, Helena Lucia and Marção, Maycon and Ruiz, Max and Courtois, Elise and Flynn, William F and Sivajothi, Santhosh and Soja, Emily and Caratù, Ginevra and Mora-Roldan, German Atzin and Dredge, B Kate and Liu, Yutian and Chasteen, Hannah and Mohenska, Monika and Nieto, Juan C and Yip, Raymond K H and Mishi, Ruvimbo D and Polo, José M and Abdalfttah, Mohmed and Sullivan, Adrienne E and Plummer, Jasmine T and Heyn, Holger and Martelotto, Luciano G. STAMP: Single-cell transcriptomics analysis and multimodal profiling through imaging. *Cell* **188**, 5100–5117.e26 (2025).
14. Nam, Anna S and Dusaj, Neville and Izzo, Franco and Murali, Rekha and Myers, Robert M and Mouhieddine, Tarek H and Sotelo, Jesus and Benbarche, Salima and Waarts, Michael and Gaiti, Federico and Tahri, Sabrin and Levine, Ross and Abdel-Wahab, Omar and Godley, Lucy A and Chaligne, Ronan and Ghobrial, Irene and Landau, Dan A. Single-cell multi-omics of human clonal hematopoiesis reveals that DNMT3A R882 mutations perturb early progenitor states through selective hypomethylation. *Nat. Genet.* **54**, 1514–1526 (2022).
15. Nam, Anna S and Kim, Kyu-Tae and Chaligne, Ronan and Izzo, Franco and Ang, Chelston and Taylor, Justin and Myers, Robert M and Abu-Zeinah, Ghaith and Brand, Ryan and Omans, Nathaniel D and Alonso, Alicia and Sheridan, Caroline and Mariani, Marisa and Dai, Xiaoguang and Harrington, Eoghan and Pastore, Alessandro and Cubillos-Ruiz, Juan R and Tam, Wayne and Hoffman, Ronald and Rabadan, Raul and Scandura, Joseph M and Abdel-Wahab, Omar and Smibert, Peter and Landau, Dan A. Somatic mutations and cell identity linked by Genotyping of Transcriptomes. *Nature* **571**, 355–360 (2019).
